# Silencing basal forebrain cholinergic input to the infralimbic cortex renders fear extinction resistant to renewal

**DOI:** 10.64898/2026.07.06.736678

**Authors:** Belinda P. P. Lay, Billy C. Chieng, Stephen Maren, Vincent Laurent

**Affiliations:** Decision Neuroscience Laboratory, School of Psychology, University of New South Wales, Sydney, NSW, Australia.; Beckman Institute for Advanced Science and Technology and Department of Psychology, University of Illinois Urbana-Champaign, Urbana, IL, USA

## Abstract

The infralimbic cortex (IL) is critical for the extinction of conditioned fear and receives dense cholinergic innervation from the basal forebrain, particularly from the horizontal limb of the diagonal band of Broca (HDB). Acetylcholine release in the IL regulates fear-related processes, but the mechanisms involved remain poorly understood. We investigated whether cholinergic projections from the HDB to the IL regulate the formation, extinction, and renewal of fear memories. Using optogenetics in male and female transgenic rats, we silenced the HDB to IL cholinergic pathway during either fear conditioning or extinction. Silencing during fear conditioning had no effect, whereas silencing during extinction enhanced extinction retrieval and prevented fear renewal. These effects were anatomically specific, as silencing the neighboring HDB to prelimbic cortex pathway had no such consequence. In ex vivo IL slices, stimulation of HDB cholinergic terminals preferentially excited interneurons over pyramidal neurons, consistent with feedforward inhibition of IL output. Finally, blockade of IL nicotinic acetylcholine receptors during extinction fully reproduced the effects of silencing the pathway, whereas muscarinic blockade attenuated renewal without affecting extinction retrieval. These findings indicate that cholinergic input from the HDB to the IL controls the acquisition and contextual specificity of extinction and identify nicotinic signaling in the IL as a potential target for improving extinction-based therapies.

**Significance Statement:** A hallmark of fear extinction is its vulnerability to disruption when environmental conditions change, leading to the return of fear outside the extinction context. This study identifies a basal forebrain cholinergic projection to the infralimbic cortex that governs this contextual control, determining whether extinction is expressed beyond the context in which it was learned. These findings are significant because they show that context specificity of extinction is not fixed but is actively regulated by a discrete cholinergic circuitry, and that removing nicotinic signaling within this circuit renders extinction resistant to relapse. This work therefore identifies nicotinic receptors in the infralimbic cortex as target for improving the efficacy of extinction-based therapies.

---

Traumatic events can have profound and lasting effects on the brain and behavior, and these effects often become maladaptive. Clinical interventions such as exposure therapy are grounded in the principles of Pavlovian conditioning, seeking to reduce maladaptive fear through repeated, safe exposure to the feared stimuli. Accordingly, decades of research have sought to unravel the mechanisms governing the formation of learned fear and, critically, its reduction (Myers and Davis, 2002; Myers and Davis, 2007; Hofmann, 2008; Quirk and Mueller, 2008; Graham and Milad, 2011; Golkar and Öhman, 2012; Delamater and Westbrook, 2014; Tovote et al., 2015; Bouton et al., 2021). A central focus has been the extinction of Pavlovian fear conditioning, in which a conditioned stimulus that signals an aversive event (e.g., a tone that had been paired with shock) is presented repeatedly in the absence of that event (the tone presented alone), leading to a reduction in fear. Although extinction reduces fear rapidly, its effects are often transient and context-specific, and fear frequently relapses (Bouton et al., 2006; Vansteenwegen et al., 2006; Laurent and Westbrook, 2009). One such instance is renewal, in which fear returns when the extinguished stimulus is presented outside of its extinction context (Bouton and Bolles, 1979; Harris et al., 2000; Bouton, 2002; Maren et al., 2013). These findings indicate that extinction does not erase the original fear memory; rather, the original fear memory and the extinction memory compete for behavioral control.

The infralimbic cortex (IL) plays a pivotal role in the extinction of conditioned fear memories. Disrupting IL function during extinction training (Morgan and LeDoux, 1995; Sierra-Mercado et al., 2006; Laurent and Westbrook, 2009; Lingawi et al., 2017b; Lay et al., 2020) or immediately after training (Burgos-Robles et al., 2007; Sotres-Bayon et al., 2009; Awad et al., 2015) impairs the formation of extinction memories and, consequently, their subsequent retrieval. Conversely, stimulating IL activity during extinction training facilitates extinction learning and strengthens later retrieval (Milad and Quirk, 2002; Vidal-Gonzalez et al., 2006; Do-Monte et al., 2015; Lingawi et al., 2017b; Lingawi et al., 2026). Such stimulation can also reduce forms of relapse, including spontaneous recovery (Wills et al., 2025) and renewal (Thompson et al., 2010). The IL, and the medial prefrontal cortex (mPFC) more broadly, receives dense cholinergic input from the basal forebrain, particularly from the horizontal limb of the diagonal band of Broca (HDB; Bloem et al., 2014b; Crimmins et al., 2023). Acetylcholine (ACh) release in the mPFC is thought to regulate a range of functions, including attention and fear extinction (Parikh et al., 2007; Santini et al., 2012; Bloem et al., 2014a; Robinson-Drummer et al., 2017; Wilson and Fadel, 2017; Fernandes-Henriques et al., 2025; Tu et al., 2025). For example, blocking muscarinic ACh receptors in the IL impairs the consolidation of fear extinction memories (Santini et al., 2012). Moreover, silencing IL projections to a subregion of the basal forebrain during extinction learning impairs within-session fear reduction without affecting extinction retrieval (Fernandes-Henriques et al., 2025). However, the role of the cholinergic input from the HDB to the IL in extinction and, importantly, in the return of extinguished fear remains largely unexplored.

The present experiments examined how cholinergic projections from the HDB to the IL regulate the formation, extinction and renewal of fear memories. We used optogenetics in transgenic rats to silence the HDB to IL cholinergic pathway (^ACh^HDB→IL) during either fear conditioning or fear extinction. The long-term consequences of this manipulation were assessed during a post-extinction test and a pair of context tests, one in the extinction context and one outside it, conducted in either the extinction and conditioning contexts (ABA renewal) or the extinction context and a second, familiar context (AAB renewal). In doing so, we hoped to offer insight into how the cholinergic system regulates fear and into the therapeutic potential of cholinergic compounds for disorders of fear and anxiety.

## Materials and Methods

### Subjects

Subjects were experimentally naïve male and female ChAT::Cre^+^ transgenic rats (*n* = 160; Rat Resource & Research Center, USA; #00658; Witten et al., 2011; Crimmins et al., 2023) or wild-type rats (*n* = 57) bred on a Long-Evans background. The rats were obtained from the Breeding Facility at the University of New South Wales (Sydney, Australia) and were at least 8 weeks old at the start of the experiments. They were housed in plastic boxes (3-4 rats per box) located in a climate-controlled colony room maintained on a 12 h light/dark cycle (lights on between 7:00 A.M. and 7:00 P.M.). Histological exclusions prevented the inclusion of an equal number of females and males in each experimental group. Nevertheless, separate statistical analyses revealed no sex differences during the various experimental stages (smallest *p* = 0.11). The Animal Ethics Committee at the University of New South Wales approved all experimental procedures, which took place during the light cycle.

### Viruses and Surgery

The following adeno-associated viruses (AAVs) were used: AAV5-EF1a-DIO-eNpHR3.0- eYFP (eNpHR3.0; Addgene; #26966) and AAV5-EF1a-DIO-eYFP (Addgene; #27056; eYFP).

At the time of surgery, male rats weighed between 250 and 420 g whereas female rats weighed between 200 and 320 g. A continuous flow of mixed isoflurane and oxygen gas was used to anesthetize rats, which were then placed in a stereotaxic frame (Kopf Instruments; California, USA). An incision was made to expose the scalp, and the incisor bar was adjusted to align bregma and lambda on the same horizontal plane. For viral infusions, holes were drilled bilaterally into the skull above the HDB at the following coordinates (indicated in mm relative to bregma; male/female when necessary): +0.9 antero-posterior (AP), ±0.65 medial-lateral (ML), -8.45/8.37 dorsal-ventral (DV) from bregma. Infusions were conducted using a 1 µl Hamilton syringe attached to an infusion pump (Pump 11 Elite Nanomite, Harvard Apparatus). A total of 0.5 µl of the appropriate AAV was infused in each hemisphere at a rate of 0.1 µl/min. The syringe was left in place for an additional 7 min to allow for diffusion. For the fiber optic implants (Doric Lenses, Quebec, Canada; MFC_400/430-0.48_10mm_RM2_FLT or RWD, Guangdong, China; R-FOC-BF400C-39NA-Fiber Optic Cannulae, Black, with 2.5 mm Ceramic Ferrule, 400 µm Core, 0.39 NA), holes were drilled bilaterally into the skull above the target brain regions at the following coordinates: IL cortex: +2.5 AP, ±2 ML at a 15° angle, -4.7/4.55 DV from bregma; PL cortex: +2.5 AP, ±1.8 ML at a 15° angle, -3.3/3.15 DV from bregma. The fiber optics were secured to the skull with four jeweler’s screws and dental cement.

For IL cannula implantation, twenty-six-gauge single-guide cannulae (Plastics One) were implanted through holes drilled in both hemispheres of the skull above the IL. The tips of the guide cannulae were aimed at the IL bilaterally using the following coordinates: +2.5 mm AP, ±2 mm ML at a 15° angle (bypassing the prelimbic cortex to avoid damage), and -4.7/4.55 mm DV from bregma. The guide cannulae were secured to the skull with four jeweler’s screws and dental cement. A dummy cannula was kept in the guide at all times except during microinjections.

Animals were allowed to recover for seven days following surgery and were handled daily for five days prior to the start of the behavioral procedures.

### Apparatus

Training and testing took place in Med Associates conditioning chambers (Vermont, USA) enclosed in sound- and light-resistant shells as described previously (Crimmins et al., 2023). The floor of the chambers consisted of stainless-steel rods (3.8 mm in diameter, spaced 16 mm apart) that could be connected to a constant current generator to deliver a 0.5 s, 0.8 mA foot shock unconditioned stimulus (US). Each chamber contained a speaker connected to a sound card to generate a 3 kHz, 90 dB pure tone conditioned stimulus (CS). The CS lasted 30 s in all experiments. Each chamber was equipped with infra-red cameras to record the rat’s behavior. A computer located in another room in the laboratory controlled the delivery of stimuli via the Med-PC V Software Suite. Two distinct physical contexts (A and B) that differed in terms of visual features, odor, and floor type were used. Context A consisted of the standard bare operant chambers with a stainless grid floor and paper bedding in the tray. In contrast, Context B had a smooth black Perspex floor, laminated sheets of paper with vertical black and white stripes (2 cm) affixed to the front wall and ceiling of the chamber, no bedding in the tray, and 10% peppermint solution (Queen).

### Behavioral procedures

On day 1, rats received two pre-exposure sessions that occurred at least two and a half hours apart. One session took place in Context A and the other in Context B (counterbalanced). Each session lasted approximately 30 min and included 2 presentations of the tone stimulus with an average intertrial interval (ITI) of 8 min (range: 7 to 9 min). On day 2, rats received fear conditioning in Context A, which involved three or four pairings of the tone with the foot shock. The first trial began 2 min after the rats were placed in the context and the ITI was the same as that used for pre-exposure. On day 3, rats received fear extinction in Context B during which the tone was presented 10 times in the absence of the foot shock. The first trial began 2 min after being placed in the context and the ITI ranged from 2 to 4 min and was 3 min on average. On day 4, rats received a post-extinction test in Context B to assess whether the manipulation had any effect on extinction retrieval. This test session was identical to fear extinction except in the absence of any manipulation. On days 5 and 6, rats received two context tests. One was conducted in the context in which extinction had taken place and the other was conducted in a context in which extinction had not taken place and constituted the renewal test. For the ABA design these were context B and context A respectively, and for the AAB design context A and context B (order counterbalanced). Each test involved 5 presentations of the tone.

### Optogenetics

Optical fibers were connected to patch cords with ceramic sleeves, and the patch cords were connected to an LED driver (Doric Lenses, Quebec, Canada). During the pre-exposure sessions, rats were tethered to a patch cord, but no light was delivered. This was to ensure habituation to the procedure. For optical silencing, orange light (625 nm; continuous; at least 8 mW at the tip of the fiber optic) was delivered to cholinergic terminals expressing eNpHR3.0 in the IL or PL. The light was activated at tone onset and ended 4.5 s after tone offset (34.5 s total duration). Prior work in the laboratory demonstrated that LED illumination alone had no effect on behavior; both groups received therefore LED illumination.

### Drugs and drug infusion

The competitive nonselective muscarinic receptor antagonist scopolamine hydrobromide (Tocris) was used to pharmacologically block muscarinic receptor activity in the IL. The nonselective and noncompetitive nicotinic receptor antagonist mecamylamine hydrochloride (Sigma-Aldrich Australia; #M9020) was used to pharmacologically block nicotinic receptor activity in the IL. Scopolamine was dissolved in 0.9% (wt/vol) non-pyrogenic saline to obtain a final concentration of 20 µg/µl. Mecamylamine was dissolved in 0.9% (wt/vol) non-pyrogenic saline to obtain the following final concentrations: 1 µg/µl and 10 µg/µl. The concentrations selected were based on prior studies (Zarrindast et al., 2005; Rezayof et al., 2006; Santini et al., 2012; Jiang et al., 2016). Saline was used as a vehicle solution. A total volume of 0.3 µl was delivered to each hemisphere at a rate of 0.1 µl/min.

Prior to fear extinction, rats were allocated to drug or vehicle groups, matched on their freezing performance during fear acquisition. Ten minutes before the extinction session, scopolamine, mecamylamine or vehicle was infused bilaterally into the IL by inserting a 33-gauge injector cannula into each guide cannula. The injector cannulae were connected to a 25 µl glass Hamilton syringe attached to an infusion pump (Harvard Apparatus). The injector cannula projected an additional 1 mm ventral to the tip of the guide cannula. The injector cannula remained in place for an additional 1 min after the infusion to allow for drug diffusion before its complete removal. Immediately after the infusion, the injector was replaced with the original dummy cannula. One day before infusions, all rats were familiarized with this procedure to minimize stress on the infusion day.

### Electrophysiology

#### Brain slice preparation

The adeno-associated virus AAV-DIO-ChR2-eYFP (ChR2) was bilaterally injected into the HDB of female ChAT::Cre+ rats 4 weeks prior to the brain slice experiment, using the coordinates mentioned above. Rats were euthanized under deep anesthesia (isoflurane 4% in air). The brain was rapidly removed and cut using a vibratome (Leica Microsystems VT1200S) in ice-cold oxygenated sucrose buffer containing (in mM): 241 sucrose, 28 NaHCO3, 11 glucose, 1.4 NaH2PO4, 3.3 KCl, 0.2 CaCl2, 7 MgCl2. Coronal brain slices (300 µm thick) were sampled and maintained at 33 °C in a submerged chamber containing physiological saline with composition (in mM): 124 NaCl, 2.5 KCl, 1.25 NaH2PO4, 1 MgCl2, 1 CaCl2, 10 glucose and 26 NaHCO3 equilibrated with 95% O2 and 5% CO2.

#### Electrophysiological recording

After equilibration for 1 h, slices were transferred to a recording chamber and neurons visualized under an upright microscope (BX50WI, Olympus) using differential interference contrast (DIC) Dodt tube optics and eYFP fluorescence, and superfused continuously (1.5 ml/min) with oxygenated physiological saline at 33 °C. Cell-attached and whole-cell patch-clamp recordings were made using electrodes (2–5 MΩ) containing internal solution (in mM): 115 K gluconate, 20 NaCl, 1 MgCl2, 10 HEPES, 11 EGTA, 5 Mg-ATP, 0.33 Na-GTP, and 5 phosphocreatine di(tris) salt (#P1937, Sigma-Aldrich), pH 7.3, osmolarity 285-290 mOsm/L. Biocytin (0.1%; #B4261, Sigma) was added to the internal solution for marking the sampled neurons during recording. Data acquisition was performed with a Multiclamp 700B amplifier (Molecular Devices), connected to a PC and interface ITC-18 (Instrutech). Liquid junction potentials of - 10 mV were not corrected. In cell-attached and then whole-cell current-clamp modes, membrane currents and potentials were sampled at 5 kHz (low pass filter 2 kHz, Axograph X, Axograph).

#### LED stimulation

Transduced ChR2 HDB terminals in the IL were stimulated either by a single LED light pulse (470 nm, 5 ms pulse, 3.5 mW, ThorLabs, Newton, NJ, USA) once per minute, or by a train of LED pulses (470 nm, 5 ms pulse, 10 Hz, 15 pulses, 3.5 mW) once every 30 s, illuminated onto the brain slice under the 40X water-immersion objective. The experiment was performed under current-clamp at resting membrane potential.

#### Post hoc histological analysis

Immediately after physiological recording, brain slices containing biocytin-filled neurons were fixed overnight in 4% paraformaldehyde/0.16 M phosphate buffer (PB) solution, rinsed, and then placed in 0.5% Triton X-100/PB for 3 days to permeabilize cells. Slices were then incubated in primary antibody for 3 days, chicken anti-GFP (GFP-1020, 1:1000, Aves Labs) to enhance eYFP signal, plus 2% horse serum and 0.2% Triton X-100/PB at 4 °C. After a rinse, this was followed by a one-step overnight incubation at 4 °C in a fluorescent avidin plus a secondary antibody: ExtrAvidin Cy3 (E4142, 1:1000, Sigma), and donkey anti-chicken Alexa 488 conjugated IgY (703-545-155, 1:1000, Jackson ImmunoResearch Inc). Stained slices were rinsed, mounted onto glass slides, dried, and coverslipped with Fluoromount-G mounting medium (00-4958-02, Invitrogen). A 2D projection was later obtained from a collated image stack using confocal laser scanning microscopy (Fluoview FV1000, BX61WI microscope, Olympus).

### Histology

Subsequent to behavioral testing, rats received a lethal dose of sodium pentobarbital (300 mg/kg; Virbac Pty. Ltd., Sydney, Australia) diluted 1:1 with 0.9% sodium chloride and were transcardially perfused with cold 4% paraformaldehyde in 0.1 M sodium phosphate buffer (PB; pH 7.5). Brains were extracted and post-fixed in the same solution at 4 °C overnight. Coronal 50 µm thick sections were cut through the mPFC and HDB with a vibratome (VT1000, Leica Microsystems; Sydney, Australia) and stored at -30 °C in a solution containing 30% ethylene glycol, 30% glycerol, and 0.1 M sodium phosphate buffer.

To achieve immunodetection of cholinergic neurons, individual free-floating sections were rinsed three times for 10 min in Tris-buffered saline (TBS: 0.25 M Tris, 0.5 M NaCl, pH 7.5). After a 40 min incubation in 0.2% Triton X-100 in TBS, sections were rinsed three times in TBS. The slices were then incubated overnight at 4 °C in TBS containing the goat anti-ChAT primary antibody (1:300; Millipore; # AB144P). The following day, the slices were rinsed three times in TBS and incubated for 60 min at room temperature with donkey anti-goat Cy3 (1:400; Jackson ImmunoResearch Laboratories; #705-165-147). After this incubation, sections were rinsed three times in TBS, mounted on Superfrost Plus-coated slides, and left to dry for 10 min before being coverslipped in Vectashield mounting medium (Vector Laboratories, # VEH- 1000). The mounting procedure just described was used to examine the spread of the viral infusions and placement of the fiber optics. Rats with inadequate viral infusion spread or misplacement of the fiber optics in the target regions, as defined by the atlas of Paxinos and Watson(Paxinos and Watson, 1997), were excluded from the statistical analyses.

Fluorescent brain sections were imaged using a spinning disk confocal system equipped with the Diskovery multi-modal imaging platform (SAR/Andor Technology) and a Zyla 4.2 sCMOS camera (Andor Technology), which allows fast capture of large mosaic images and high-sensitivity, high-dynamic range confocal imaging. The Diskovery platform was mounted on a Nikon Eclipse TiE microscope body with a motorized stage and the Nikon Perfect Focus System, and image acquisition was controlled by Nikon NIS-Elements software. All images were acquired with a Nikon 20X objective (0.75 N.A. CFI Plan Apo). All images were then processed using Open Source ImageJ/Fiji software(Rueden et al., 2017). Viral expression specificity (Figure 1J) was assessed through manual cell counting and involved calculating the percentage of fluorophore-labeled (eYFP) cells relative to the total number of ChAT-positive cells, and a percentage of ChAT-positive cells relative to the total number of fluorophore-labeled (eYFP) cells. The total number of positive cells was quantified by sampling the HDB region from two to three sections per rat. Each rat represented a single observation.

**Fig. 1.**
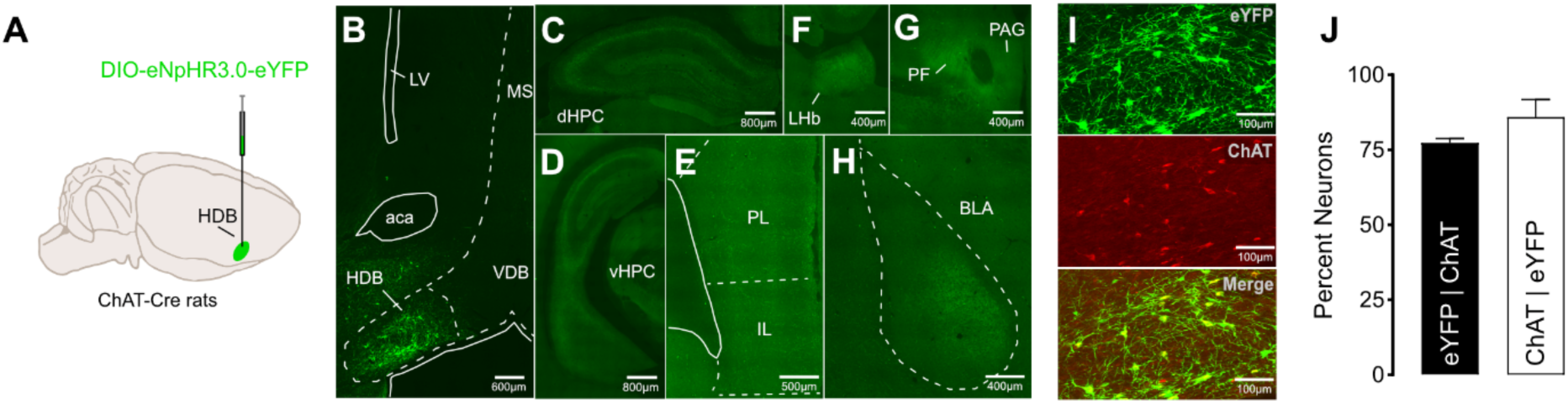
*HDB cholinergic neurons are selectively transduced and innervate the mPFC* **A.** ChAT::Cre+ rats were bilaterally infused in the HDB with DIO-eNpHR3.0-eYFP. **B.** Micrographs showing HDB viral expression. **C-H.** HDB cholinergic neurons project to the dorsal (dHPC) and ventral (vHPC) hippocampus, the medial prefrontal cortex (PL and IL), lateral habenula (LHb), the periaqueductal gray (PAG), the parafascicular thalamic nucleus (PF), and the basolateral amygdala (BLA). **I.** Individual channel and composite micrographs showing HDB viral expression. Viral expression was restricted to HDB cholinergic neurons. ChAT choline acetyltransferase. **J.** Most HDB cholinergic neurons were eYFP-positive (eYFP | ChAT; n = 4 rats (3M, 1F); 80 ChAT neurons total) and most HDB eYFP-positive neurons were cholinergic (ChAT | eYFP; 87 eYFP-positive neurons total).

### Statistical analysis

Freezing was used to assess conditioned fear. It was defined as the absence of all movement except that required for breathing, accompanied by a hunched posture(Blanchard and Blanchard, 1969; Fanselow, 1980). Each rat was observed every 2 s and scored as either freezing or not freezing by two observers, one of whom was blind to group assignment. The correlation between the scores was high (Pearson r > 0.9). For each rat, freezing was expressed as the percentage of observations scored as freezing during the baseline period (2 min before the first tone presentation at each stage) and each tone presentation. The levels of freezing did not include freezing during the ITI. Data were analyzed in SPSS 25.0 (IBM, New York, USA) using repeated measures analysis of variance (ANOVA). Significance was set at α = 0.05. Where appropriate, Bonferroni adjustments were made for multiple comparisons. Baseline levels of freezing were analyzed using independent samples t-tests or a one-way ANOVA. Effects of trial, where reported, were measured with contrasts testing for the presence of a linear trend. If interactions were detected, follow-up simple effects analyses were conducted to determine their source. Data for each experiment and phase were reported using all trials unless otherwise specified. One rat from the nAChR blockade study failed to acquire fear to the tone and was excluded from the statistical analysis.

## Results

### HDB cholinergic neurons are selectively transduced and innervate the mPFC

We first characterized the viral transduction and projection targets of HDB cholinergic neurons in transgenic rats expressing Cre recombinase in choline acetyltransferase-expressing neurons (ChAT::Cre+; Witten et al., 2011). Rats were infused in the HDB (Figs. 1A, B) with a Cre- dependent adeno-associated virus encoding the neuronal silencer halorhodopsin and the fluorophore eYFP (eNpHR3.0 virus). eYFP-positive terminals were observed in the prelimbic (PL) and IL cortices, basolateral amygdala, dorsal and ventral hippocampus, lateral habenula, and lateral periaqueductal gray (Figs. 1C-H). Viral expression was largely restricted to cholinergic neurons (Figs. 1I, J). Most HDB cholinergic neurons were eYFP-positive (eYFP|ChAT; *n* = 4; 81 ChAT neurons total), and most HDB eYFP-positive neurons were cholinergic (ChAT|eYFP; 87 eYFP-positive neurons total). These results are consistent with the literature(Bloem et al., 2014b; Chaves-Coira et al., 2018; Crimmins et al., 2023) and confirm that HDB cholinergic neurons innervate the IL and PL cortices of the mPFC as well as other regions implicated in fear (Fanselow and LeDoux, 1999; Zelikowsky et al., 2012; Duvarci and Pare, 2014; Watson et al., 2016; Arico et al., 2017; Assareh et al., 2017; Lingawi et al., 2017a; Lay et al., 2018; Durieux et al., 2020; Oh and Han, 2020; Velazquez-Hernandez and Sotres-Bayon, 2021; Sachella et al., 2022; Zaki et al., 2022).

### ^ACh^HDB→IL, but not ^ACh^HDB→PL, silencing during extinction enhances extinction retrieval and prevents renewal

Having confirmed that we could selectively transduce HDB cholinergic neurons and that these neurons project to the IL, we sought to determine whether this cholinergic projection is critical for fear extinction. ChAT::Cre+ rats were infused in the HDB with the eNpHR3.0 or control eYFP virus and were implanted with fiber optics above the IL (Figs. 2B, C, S1A-C). All rats were trained to fear a tone paired with a shock in context A (Fig. 2A). Fear to the tone was then extinguished in context B. Both groups received LED illumination during extinction, such that the ^ACh^HDB→IL pathway was silenced in eNpHR3.0 rats and left intact in eYFP control rats. The long-term effects of silencing were assessed in context B during a post-extinction test conducted in the absence of illumination. To determine whether the loss of fear survived a context change, rats were then given two context tests: one in context A and one in context B (order counterbalanced). Renewal would be evident as higher fear in context A than in the extinction context B.

**Fig. 2.**
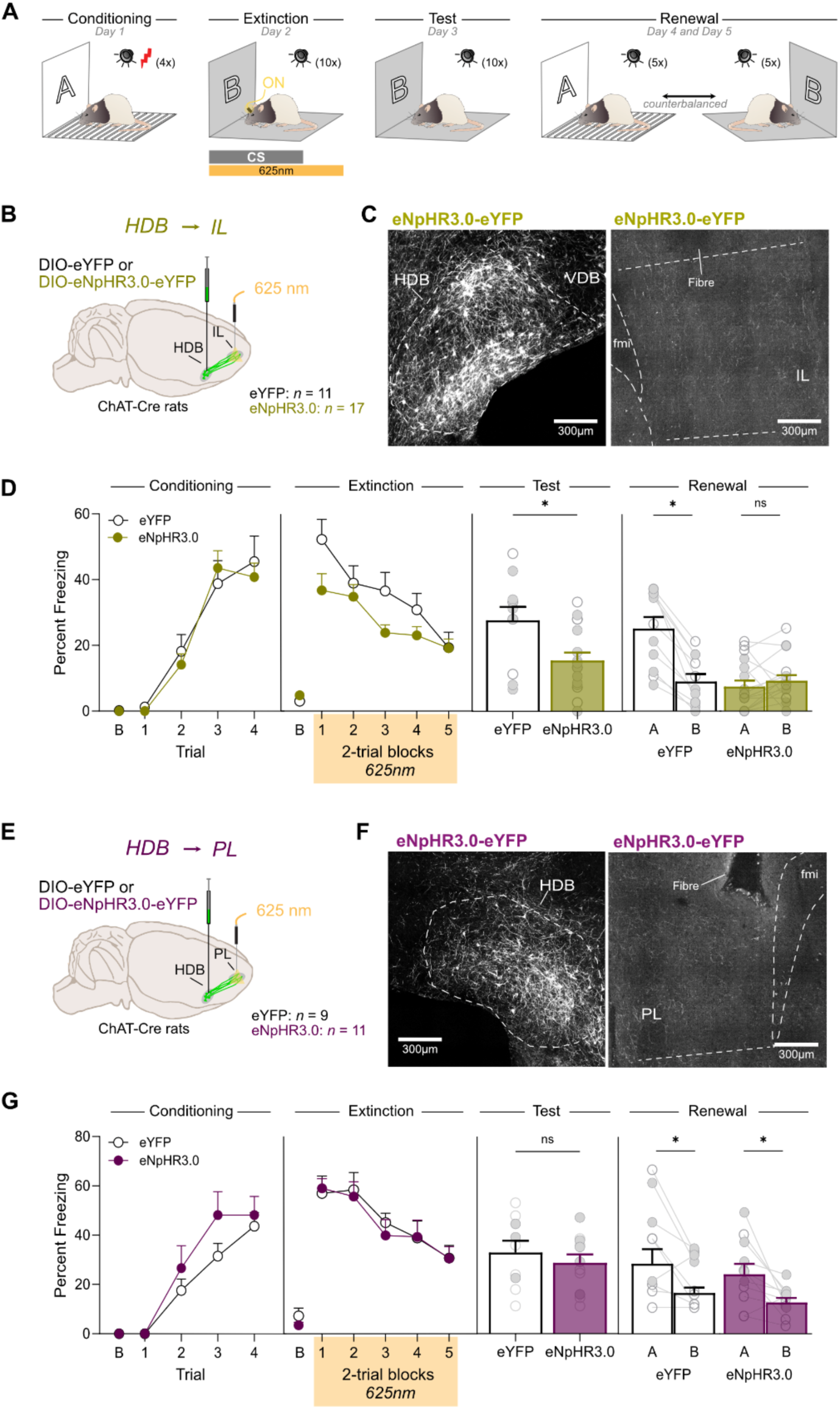
^ACh^HDB→IL, but not ^ACh^HDB→PL, silencing during extinction enhances extinction retrieval and prevents renewal **A.** Schematic representation of the experimental design. **B.** ChAT::Cre+ rats were bilaterally infused in the HDB with either DIO-eYFP (n = 4M, 7F) or DIO-eNpHR3.0-eYFP (n = 6M, 11F) and fiber optics were bilaterally implanted above the IL. **C**. Micrographs showing DIO- eNpHR3.0-eYFP expression in HDB cholinergic neurons (left), eYFP-positive IL cholinergic terminals and fiber optic placements (right). **D.** Baseline freezing (B) to the context prior to first tone CS presentation. Fear conditioning was also similar between groups. ^ACh^HDB→IL silencing reduced freezing during extinction. ^ACh^HDB→IL silencing reduced freezing during the post-extinction test. ^ACh^HDB→IL silencing abolished renewal. **E.** ChAT::Cre+ rats were bilaterally infused in the HDB with either DIO-eYFP (n = 6M, 3F) or DIO-eNpHR3.0-eYFP (n = 6M, 5F) and fiber optics were bilaterally implanted above the PL. **F.** Micrographs showing DIO-eNpHR3.0-eYFP expression in HDB cholinergic neurons (left), eYFP-positive PL cholinergic terminals and fiber optic placements (right). **G**. Baseline freezing (B) to the context prior to first tone CS presentation. Fear conditioning was similar between groups. ^ACh^HDB→PL silencing reduced freezing during extinction. ^ACh^HDB→PL silencing had no effect on freezing during the post-extinction test. ^ACh^HDB→PL silencing had no effect on fear renewal. Data are shown as mean + SEM. Asterisks denote significant effect (*p<0.05). n.s. nonsignificant. Each light gray dot corresponds to one animal.

The freezing data are presented in Figures 2D, S1D. Baseline levels of freezing during training and test sessions did not differ between the groups (max *t*(26) = 1.26, *p* = 0.22). Fear conditioning was successful, and freezing gradually increased across trials (*F*_1,26_ = 131.24, *p* < 0.001). This increase was similar across groups, as there was neither a main effect of group (*F*_1,26_ = 0.097, *p* = 0.76) nor a group by trial interaction (*F*_1,26_ = 0.003, *p* = 0.95). Extinction was also successful and freezing gradually decreased across trials (*F*_1,26_ = 39.27, *p* < 0.001). There was no main effect of group (*F*_1,26_ = 2.97, *p* = 0.097) and no group by trial interaction (*F*_1,26_ = 1.97, *p* = 0.17). During the post-extinction test, eNpHR3.0 rats froze less than eYFP control rats (*F*_1,26_ = 7.55, *p* = 0.011). Silencing also abolished fear renewal. There was a main effect of group, with silenced rats freezing less than controls overall (*F*_1,26_ = 9.20, *p* = 0.005), and a main effect of context, with freezing higher in context A than in context B (*F*_1,26_ = 22.73, *p* < 0.001). Critically, there was a group by context interaction (*F*_1,26_ = 35.63, *p* < 0.001): control rats displayed renewal, freezing more in context A than in context B (*F*_1,26_ = 47.47, *p* < 0.001), whereas eNpHR3.0 rats did not, freezing at similar levels in the two contexts (*F*_1,26_ = 0.92, *p* = 0.35).

Having found that ^ACh^HDB→IL silencing facilitates extinction retrieval and abolishes renewal, we asked whether these effects are specific to the IL. The PL is immediately adjacent to the IL, also receives HDB cholinergic inputs (Fig. 1E), and is itself known to regulate fear (Corcoran and Quirk, 2007; Laurent and Westbrook, 2009; Sierra-Mercado et al., 2011; Kim et al., 2013; Sharpe and Killcross, 2015a; Sharpe and Killcross, 2015b). We therefore examined the impact of silencing the HDB to PL cholinergic pathway (^ACh^HDB→PL) during fear extinction, using a procedure identical to that described above except that fiber optics were implanted above the PL (Figs. 2E, F, S1 E-G). The freezing data are presented in Figures 2G, S1H. Baseline levels of freezing during training and test sessions did not differ between the groups (max *t*(18) = 1.65, *p* = 0.16) except during the post-extinction test where ^ACh^HDB→PL silencing reduced contextual freezing relative to the eYFP control rats (*t*(18) = 2.67, *p* = 0.016). The source of this difference remains unclear but is unlikely to have affected overall results given it was confined to a single stage. Fear conditioning was successful and freezing gradually increased across trials (*F*_1,18_ = 102.71, *p* < 0.001). This increase was equivalent across groups, as there was neither a main effect of group (*F*_1,18_ = 2.07, *p* = 0.17) nor a group by trial interaction (*F*_1,18_ = 0.47, *p* = 0.50). Extinction was also successful and freezing gradually decreased across trials (*F*_1,18_ = 54.42, *p* < 0.001). ^ACh^HDB→PL silencing had no effect on within-session freezing during extinction (*F*_1,18_ = 0.026, *p* = 0.87) and there was no group by trial interaction (*F*_1,18_ = 0.003, *p* = 0.96). In contrast to ^ACh^HDB→IL silencing, ^ACh^HDB→PL silencing had no effect on extinction retrieval during the post-extinction test, with all rats displaying similar freezing (*F*_1,18_ = 0.52, *p* = 0.48). The silencing also had no effect on renewal. There was a main effect of context, with freezing higher in context A than in context B (*F*_1,18_ = 15.90, *p* < 0.001), but no main effect of group (*F*_1,18_ = 0.78, *p* = 0.39) and no group by context interaction (*F*_1,18_ = 0.003, *p* = 0.96). Thus, both groups renewed, freezing more in context A than in the extinction context B.

Together, these findings show that silencing the ^ACh^HDB→IL pathway during extinction, but not silencing the ^ACh^HDB→PL pathway, both facilitated retrieval of the extinction memory and prevented the renewal of freezing.

### ^ACh^HDB→IL and ^ACh^HDB→PL silencing during conditioning does not alter the formation or extinction of fear

To determine whether the ^ACh^HDB→IL and ^ACh^HDB→PL pathways regulate the formation of fear memories, we used the same approach as before (Figs. 3A-C, 3E, F, S2A-C, E-G), except that silencing occurred during conditioning. The freezing data for ^ACh^HDB→IL are presented in Figs. 3D, S2D. Baseline levels of freezing during training and test sessions did not differ between the groups (max *t*(25) = 1.17, *p* =0.25). Fear conditioning was successful and freezing gradually increased across trials (*F*_1,25_ = 105.37, *p* < 0.001). This increase was similar across groups, as there was neither a main effect of group (*F*_1,25_ = 0.11, *p* = 0.74) nor a group by trial interaction (*F*_1,25_ = 2.23, *p* = 0.15). Extinction was also successful and freezing gradually decreased across trials (*F*_1,25_ = 80.83, *p* < 0.001), again with no main effect of group (*F*_1,25_ = 0.27, *p* = 0.61) and no group by trial interaction (*F*_1,25_ = 1.55, *p* = 0.22). ^ACh^HDB→IL silencing during fear conditioning had no effect on extinction retrieval during the post-extinction test with all rats displaying similar freezing (*F*_1,25_ = 1.46, *p* = 0.24). The silencing also had no effect on the renewal of fear. There was a main effect of context, with higher freezing in context A than in context B (*F*_1,25_ = 45.23, *p* < 0.001) but no main effect of group (*F*_1,25_ = 0.38, *p* = 0.54) and no group by context interaction (*F*_1,25_ = 1.33, *p* = 0.26).

**Fig. 3.**
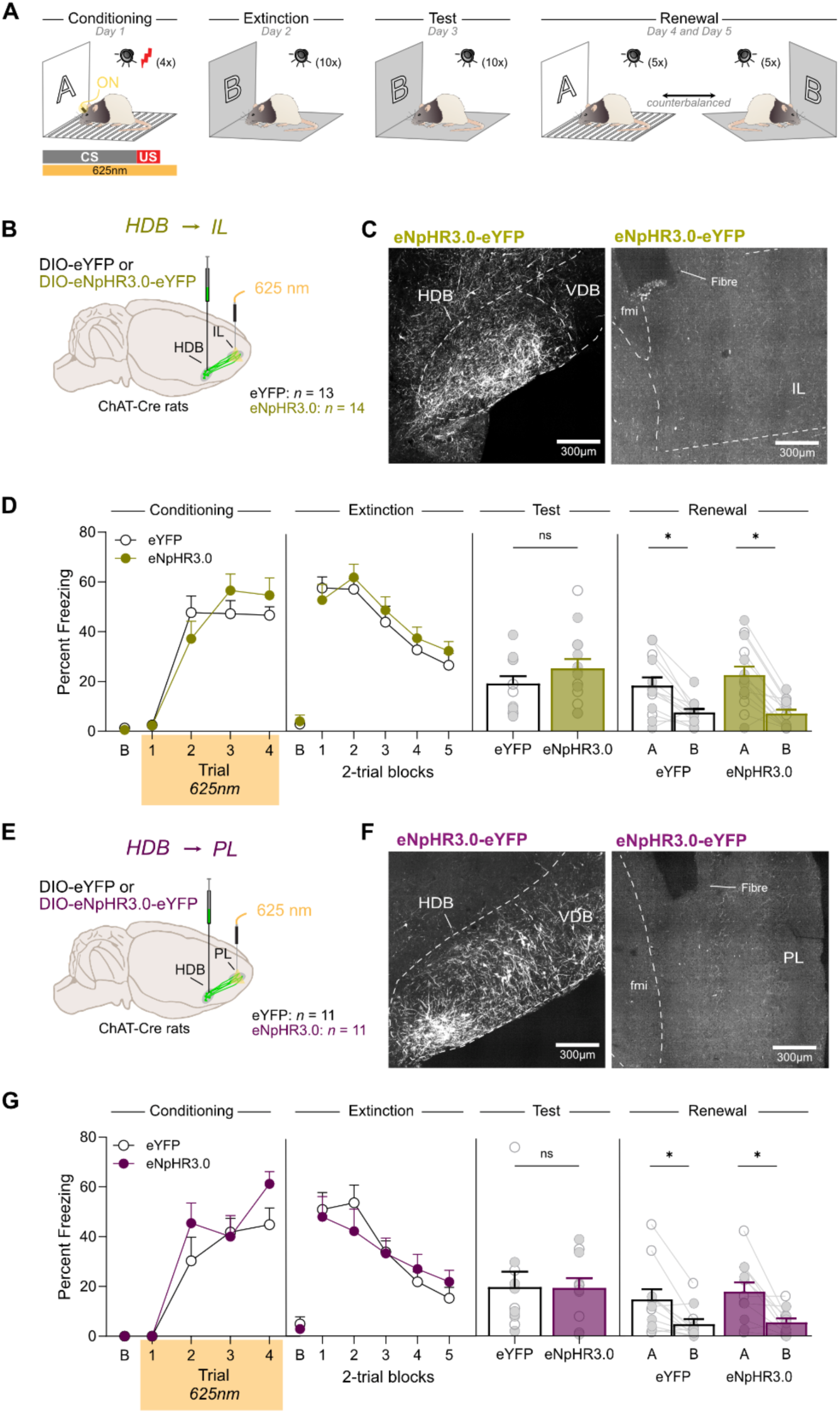
^ACh^HDB→IL and ^ACh^HDB→PL silencing during conditioning does not alter the formation or extinction of fear **A.** Schematic representation of the experimental design. **B.** ChAT::Cre+ rats were bilaterally infused in the HDB with either DIO-eYFP (n = 5M, 8F) or DIO-eNpHR3.0-eYFP (n = 6M, 8F) and fiber optics were bilaterally implanted above the IL. **C**. Micrographs showing DIO- eNpHR3.0-eYFP expression in HDB cholinergic neurons (left), eYFP-positive IL cholinergic terminals and fiber optic placements (right). **D.** Baseline freezing (B) to the context prior to first tone CS presentation. Fear conditioning was similar between groups. ^ACh^HDB→IL silencing during conditioning had no effect on within-session freezing during extinction. ^ACh^HDB→IL silencing during fear conditioning had no effect during the post-extinction test. The silencing also had no effect on the renewal of fear. **E.** ChAT::Cre+ rats were bilaterally infused in the HDB with either DIO-eYFP (n = 4M, 7F) or DIO-eNpHR3.0-eYFP (n = 4M, 7F) and fiber optics were bilaterally implanted above the PL. **F.** Micrographs showing DIO- eNpHR3.0-eYFP expression in HDB cholinergic neurons (left), eYFP-positive PL cholinergic terminals and fiber optic placements (right). **G**. Baseline freezing (B) to the context prior to first tone CS presentation. Fear conditioning was similar between groups. ^ACh^HDB→PL silencing during fear conditioning had no effect on freezing during extinction ^ACh^HDB→PL silencing had no effect during the post-extinction test. The silencing had also no effect on renewal. Data are shown as mean + SEM. Asterisks denote significant effect (*p<0.05). n.s. nonsignificant. Each light gray dot corresponds to one animal.

The freezing data for ^ACh^HDB→PL are presented in Figs. 3G, S2H. Baseline levels of freezing during training and test sessions did not differ between the groups (max *t*(20) = 1.44, *p* = 0.17). Fear conditioning was successful and freezing gradually increased across trials (*F*_1,20_ = 110.86, *p* < 0.001). This increase was similar across groups, as there was neither a main effect of group (*F*_1,20_ = 1.48, *p* = 0.24) nor a group by trial interaction (*F*_1,20_ = 1.09, *p* = 0.31). Extinction was also successful and freezing gradually decreased across trials (*F*_1,20_ = 44.55, *p* < 0.001), again with no main effect of group (*F*_1,20_ = 0.008, *p* = 0.93) and no group by trial interaction (*F*_1,20_ = 1.95, *p* = 0.18). ^ACh^HDB→PL silencing during fear conditioning had no effect on extinction retrieval during the post-extinction test with all rats displaying similar freezing (*F*_1,20_ = 0.002, *p* = 0.97). The silencing had also no effect on renewal. There was a main effect of context, with freezing higher in context A than in context B (*F*_1,20_ = 22.84, *p* < 0.001), but no main effect of group (*F*_1,20_ = 0.28, *p* = 0.60) and no group by context interaction (*F*_1,20_ = 0.26, *p* = 0.61). Together, these results indicate that neither the ^ACh^HDB→IL nor the ^ACh^HDB→PL pathway is required for the formation of fear memories.

### ^ACh^HDB→IL silencing during extinction generalizes the loss of fear to familiar contexts

We next asked whether ^ACh^HDB→IL silencing produces a more generalized extinction memory that suppresses conditioned freezing even outside the extinction context. We therefore used an AAB design, in which conditioning, extinction and the post-extinction test occurred in context A and freezing was then assessed in either that context or a distinct, familiar context B (Fig. 4A). The freezing data are presented in Figs. 4D, S3D. Baseline levels of freezing during training and test sessions did not differ between the groups (max *t*(19) = 1.32, *p* = 0.20). Fear conditioning was successful and freezing gradually increased across trials (*F*_1,19_ = 94.67, *p* < 0.001). This increase was similar across groups, as there was neither a main effect of group (*F*_1,19_ = 2.32, *p* = 0.14) nor a group by trial interaction (*F*_1,19_ = 0.13, *p* = 0.73). Extinction was also successful and freezing gradually decreased across trials (*F*_1,19_ = 73.44, *p* < 0.001), with no group by trial interaction (*F*_1,19_ = 0.15, *p* = 0.70). However, ^ACh^HDB→IL silencing reduced within-session freezing during extinction (*F*_1,19_ = 6.47, *p* = 0.02). ^ACh^HDB→IL silencing also reduced freezing during the post-extinction test (*F*_1,19_ = 10.59, *p* = 0.004) and also abolished fear renewal. There was a main effect of group, with silenced rats freezing less than controls overall (*F*_1,19_ = 14.88, *p* < 0.001), and a main effect of context, with freezing higher in context B than in context A (*F*_1,19_ = 9.88, *p* = 0.005). Critically, there was a group by context interaction (*F*_1,19_ = 7.88, *p* = 0.011): control rats displayed renewal, freezing more in context B than in context A (*F*_1,19_ = 18.59, *p* < 0.001), whereas rats with ^ACh^HDB→IL silencing froze at similar levels in both contexts (*F*_1,19_ = 0.054, *p* = 0.82).

**Fig. 4.**
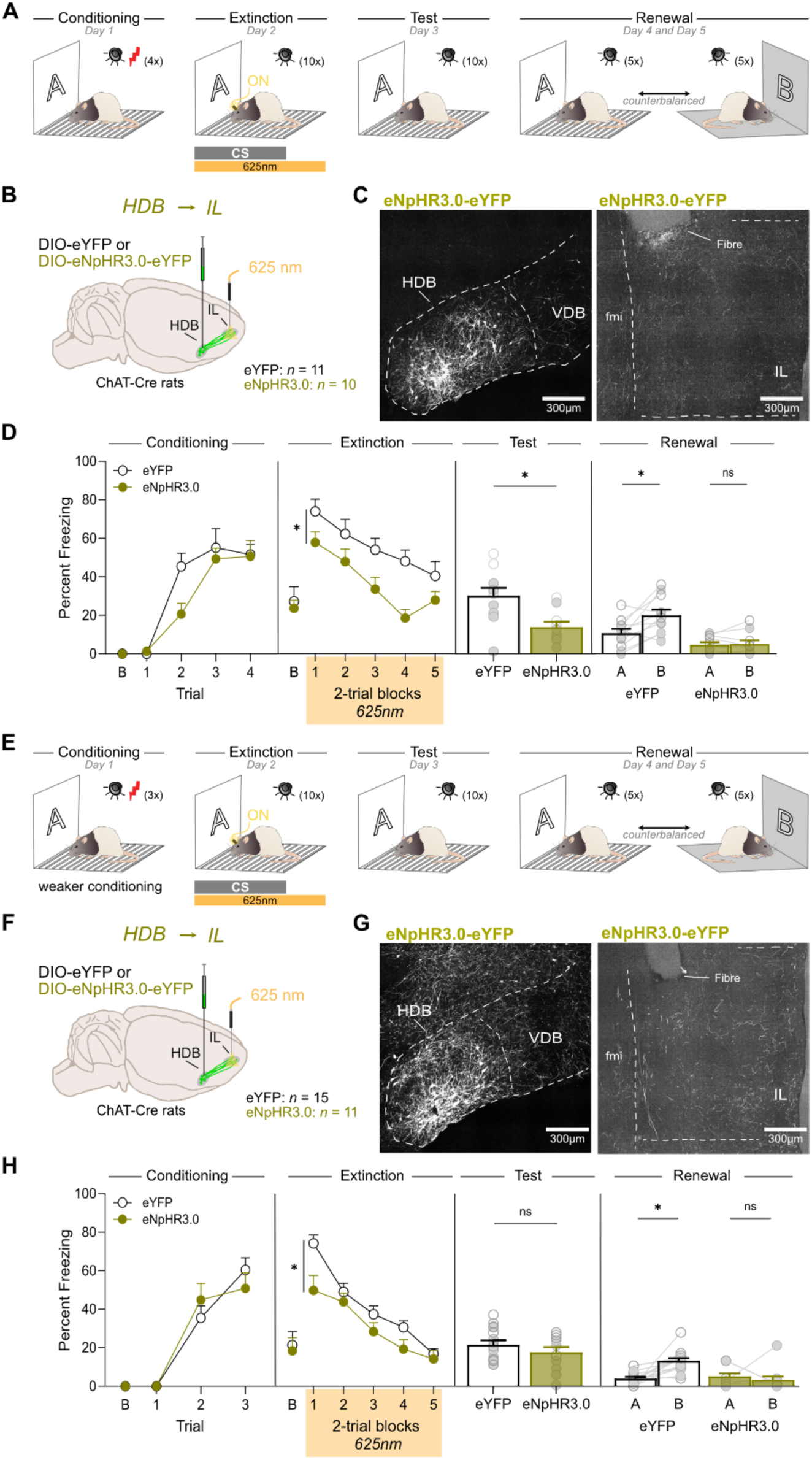
^ACh^HDB→IL silencing during extinction generalizes the loss of fear to familiar contexts **A.** Schematic representation of the experimental design. **B.** ChAT::Cre+ rats were bilaterally infused in the HDB with either DIO-eYFP (n = 5M, 6F) or DIO-eNpHR3.0-eYFP (n = 5M, 5F) and fiber optics were bilaterally implanted above the IL. **C**. Micrographs showing DIO- eNpHR3.0-eYFP expression in HDB cholinergic neurons (left), eYFP-positive IL cholinergic terminals and fiber optic placements (right). **D.** Baseline freezing (B) to the context prior to first tone CS presentation. Fear conditioning was also similar between groups. ^ACh^HDB→IL silencing reduced freezing during extinction. ^ACh^HDB→IL silencing also reduced freezing during the post-extinction test. The silencing also abolished fear renewal. **E.** ChAT::Cre+ rats were bilaterally infused in the HDB with either DIO-eYFP (n = 9M, 6F) or DIO-eNpHR3.0- eYFP (n = 4M, 7F) and fiber optics were bilaterally implanted above the IL. **F.** Micrographs showing DIO-eNpHR3.0-eYFP expression in HDB cholinergic neurons (left), eYFP-positive IL cholinergic terminals and fiber optic placements (right). **G**. Baseline freezing (B) to the context prior to first tone CS presentation. Fear conditioning was also similar between groups. ^ACh^HDB→IL silencing reduced freezing during extinction. ^ACh^HDB→IL silencing had no effect on freezing during the post-extinction test. ^ACh^HDB→IL silencing abolished fear renewal. Data are shown as mean + SEM. Asterisks denote significant effect (*p<0.05). n.s. nonsignificant. Each light gray dot corresponds to one animal.

The next experiment tested whether the loss of renewal following ^ACh^HDB→IL silencing in our previous experiments simply reflects a floor effect, as freezing was already very low at the post-extinction test, leaving little room for an increase that could reveal renewal. To address this, we sought to reduce freezing in the control group to generate similar performance in the two groups before the context tests. Our approach involved reducing the number of tone-shock pairings during conditioning (Fig. 4E). The freezing data are presented in Figs. 4H, S3H. Baseline levels of freezing during training and test sessions did not differ between the groups (max *t*(24) = 0.83, *p* = 0.42). Fear conditioning was successful and freezing gradually increased across trials (*F*_1,24_ = 122.29, *p* < 0.001). This increase was similar across groups, as there was neither a main effect of group (*F*_1,24_ = 0.00, *p* = 0.99) nor a group by trial interaction (*F*_1,24_ = 0.90, *p* = 0.35). Extinction was also successful and freezing gradually decreased across trials (*F*_1,24_ = 105.38, *p* < 0.001) with no group by trial interaction (*F*_1,24_ = 2.81, *p* = 0.11). However, ^ACh^HDB→IL silencing reduced within-session freezing during extinction (*F*_1,24_ = 5.48, *p* = 0.028). In contrast to the previous experiments, the two groups did not differ at the post-extinction test (*F*_1,24_ = 1.27, *p* = 0.27). Yet despite comparable levels of freezing before the context tests, ^ACh^HDB→IL silencing again abolished fear renewal. There was a main effect of group, with silenced rats freezing less than controls overall (*F*_1,24_ = 10.77, *p* = 0.003), and a main effect of context, with freezing higher in context B than in context A (*F*_1,24_ = 5.55, *p* = 0.027). Critically, there was a group by context interaction (*F*_1,24_ = 12.31, *p* = 0.002): control rats displayed renewal, freezing more in context B than in context A (*F*_1,24_ = 20.32, *p* < 0.001), whereas rats with ^ACh^HDB→IL silencing froze at similar levels in both contexts (*F*_1,24_ = 0.58, *p* = 0.46).

Together, these data show that silencing cholinergic input to the IL promotes the generalization of extinction to novel contexts, even when both groups entered the context tests with comparably levels of freezing.

### HDB cholinergic input excites IL interneurons and inhibits pyramidal neurons

We found that optogenetic silencing of the ^ACh^HDB→IL pathway during fear extinction enhances extinction retrieval and prevents renewal. Acetylcholine acts through two main classes of receptors, muscarinic (mAChR) and nicotinic (nAChR) acetylcholine receptors, to influence fear learning and extinction (Wilson and Fadel, 2017). However, optogenetic manipulations in ChAT::Cre+ rats do not distinguish between these receptor classes. To address this, we used slice electrophysiology to assess how HDB-derived acetylcholine influences IL neurons and to identify which acetylcholine receptor(s) underlie these behavioral effects.

Using ChAT::Cre+ rats expressing Cre-dependent ChR2, we selectively triggered acetylcholine release from HDB terminals in ex vivo IL slices. This stimulation produced mixed responses in pyramidal neurons (Fig. 5A left panel): 3/28 were excited (2.7 ± 0.7 mV depolarization), 10 of 28 were inhibited (1.2 ± 0.1 mV hyperpolarization), and 15/28 showed no response. In contrast, non-pyramidal neurons (interneurons) showed a markedly different response profile (Fig. 5A right panel), with most cells strongly excited (16/19, 7.3 ± 1.4 mV), none inhibited (0/19), and a small proportion showing no response (3/19; χ² = 26.13, df = 2, *p* < 0.001). The inhibited pyramidal neurons were primarily located in deeper layers (L1–3: 1/7; L5/6: 9/21; Fig. 5B), whereas the excited interneurons were distributed across layers (L1–3: 12/14; L5/6: 4/5; Figs. 5C and D).

**Fig. 5.**
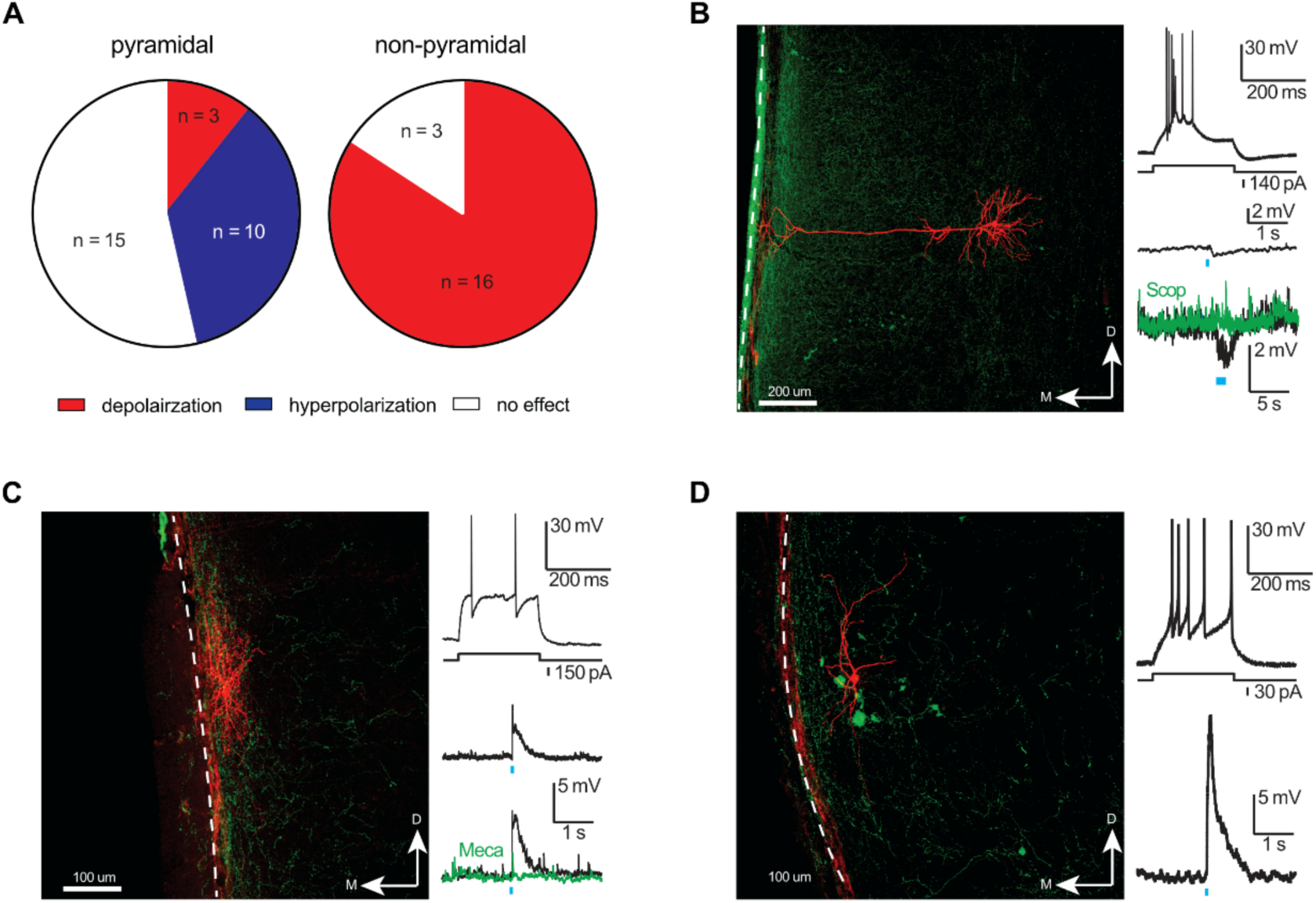
HDB cholinergic input excites IL interneurons and inhibits pyramidal neurons [**A.** Grouped data of IL neuronal response (pyramidal vs non-pyramidal) to LED stimulation with either depolarization, hyperpolarization or no effect. **B.** Morphology and physiology of a deep layer pyramidal neuron (left and top right panels). Red is biocytin and green is HDB terminal labelling in IL. Dotted white line as midline of the recorded brain slice. Example of a deep layer pyramidal neuron that is slightly hyperpolarized by LED stimulation (right middle panel). Right bottom panel shows a hyperpolarization cholinergic response blocked by scopolamine (3 uM, green trace). The LED stimulation (blue bar) was a train of 15 pulses at 10 Hz. **C.** Morphology and physiology of the Layer 1 NGF neuron (left and top right panels). Red is biocytin and green is HDB terminal labelling in IL. Dotted white line as midline of the recorded brain slice. Layer 1 NGF neuron excited by LED stimulation (blue bar, 470 nm, 5 ms pulse, 3.5 mW) with a depolarization (right middle panel). Right bottom panel shows a depolarisation cholinergic response blocked by mecamylamine (10 uM, green trace). **D.** Morphology and physiology of CA neuron (left and top right panels). Red is biocytin and green is HDB terminal labelling in IL. Dotted white line as midline of the recorded brain slice. Right bottom panel shows an example of a CA neuron in IL strongly depolarized by HDB terminal stimulation.

In the IL, stimulation of HDB cholinergic input inhibited pyramidal neurons more often than it excited them, likely via feedforward inhibition. Most of the excited interneurons were located in superficial layers (Fig. 5C right middle panel and Fig. 5D right bottom panel, respectively), where they resembled neurogliaform (NGF) and classic-accommodating (CA) subtypes (Wozny and Williams, 2011; Brombas et al., 2014; Schuman et al., 2019; Figs. 5C and 5D left and top right panels). Both populations were depolarized by HDB terminal stimulation (NGF: 3.9 ± 1.0 mV, n = 7; CA: 16.3 ± 1.9 mV, n = 3), and these responses were partially reduced by nAChR blockade (mecamylamine, 10 μM: 58.2 ± 9.5% reduction, n = 5; Fig. 5C right bottom panel). By contrast, inhibition of pyramidal neurons was abolished by mAChR antagonism (scopolamine, 3 μM, n = 6; Fig. 5B right bottom panel).

Taken together, these findings suggest that HDB cholinergic input has a predominantly inhibitory influence on IL pyramidal neurons that responded to stimulation, plausibly through feedforward inhibition. The pharmacological results point to contribution from both receptor classes, with nAChRs partly supporting the excitation of interneurons and mAChRs required for the inhibition of pyramidal neurons, although the present study does not establish how these mechanisms combine. If this influence also operates in vivo, silencing the ^ACh^HDB→IL pathway would be expected to disinhibit IL pyramidal neurons, which would be consistent with the enhanced suppression of fear that we observed after extinction.

### Nicotinic receptor blockade in the IL reproduces the effects of ^ACh^HDB→IL silencing

Our electrophysiology experiments implicated both receptor classes in the action of HDB cholinergic input on IL neurons, with nAChRs contributing to excitation of interneurons and mAChRs to the inhibition of pyramidal neurons. To determine the contribution of each class to the behavioral effects of silencing this pathway, we blocked mAChR or nAChR activity in the IL during extinction with scopolamine or mecamylamine, respectively. Rats were trained to fear a tone in context A (Figs. 6A, C, S4A, C). Ten minutes before extinction, they were assigned to a vehicle or drug group and given intra-IL infusion: scopolamine (20 µg/µl) for the mAChR study, or mecamylamine (1 or 10 µg/µl) for the nAChR study, at concentrations based on prior work (Zarrindast et al., 2005; Rezayof et al., 2006; Santini et al., 2012; Jiang et al., 2016). Fear was then extinguished in context B, and the effects of blockade were assessed drug free in context B during a post-extinction test. Rats were then given two context tests, one in context A and one in context B (order counterbalanced), to determine whether any loss of fear survived a change in context.

**Fig. 6.**
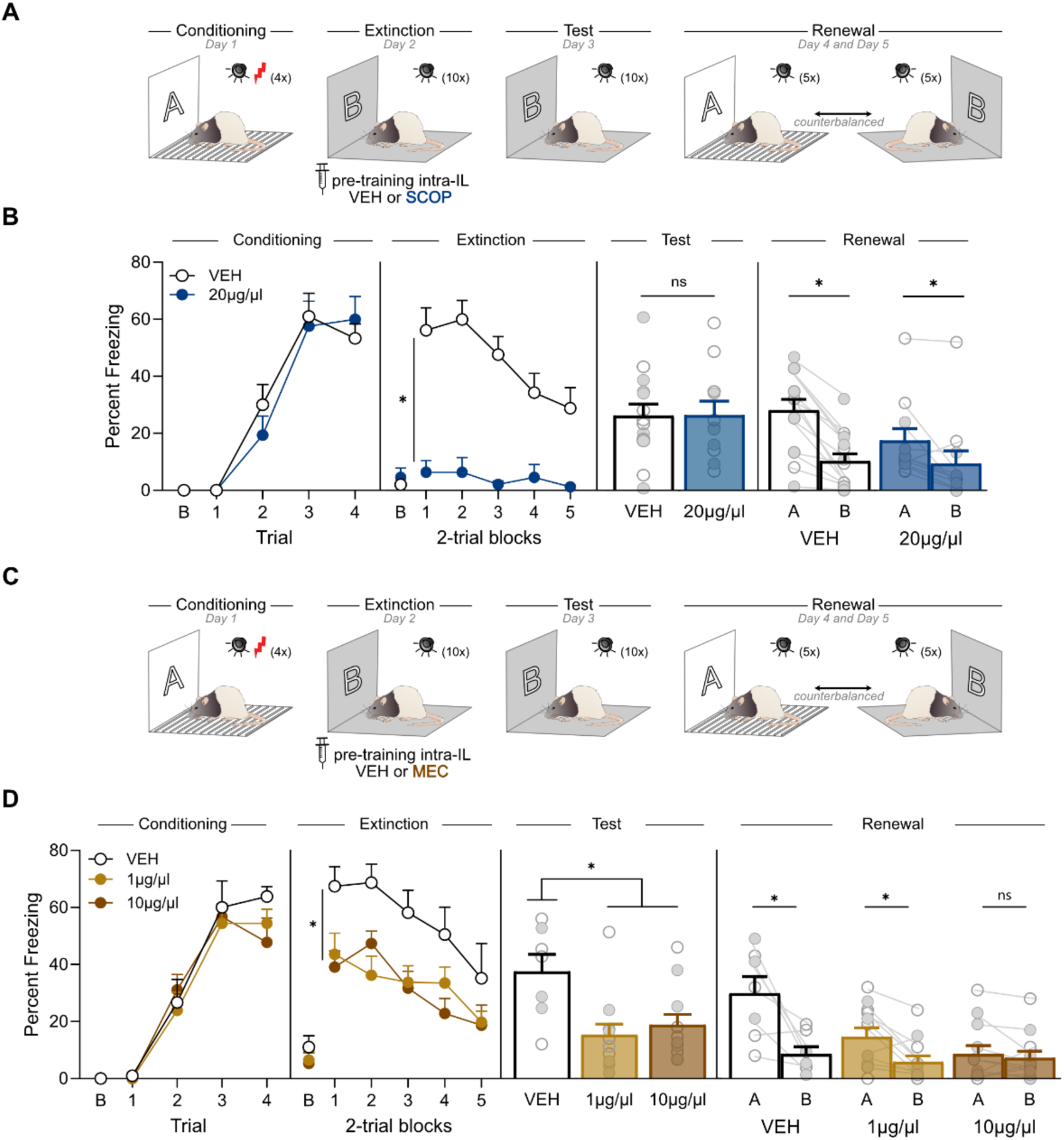
Nicotinic receptor blockade in the IL reproduces the effects of ^ACh^HDB→IL silencing **A.** Schematic representation of the experimental design (*n*s: VEH = 5M, 9F; 20µg/µl = 6M, 5F). **B.** Baseline freezing (B) to the context prior to first tone CS presentation. Fear conditioning was also similar between groups. Blocking mAChR with scopolamine reduced freezing during extinction. mAChR blockade had no effect on freezing during the post­extinction test. mAChR blockade had no effect on fear renewal. **C.** Schematic representation of the experimental design (*n*s: VEH = 4M, 3F; 1µg/µl = 6M, 6F; 10µg/µl = 6M, 6F). **D.** Baseline freezing (B) to the context prior to first tone CS presentation. Fear conditioning was also similar between groups. nAChR blockade reduced freezing during extinction. nAChR blockade reduced freezing during the post-extinction test. nAChR blockade silencing abolished fear renewal in a dose-dependent manner. Data are shown as mean + SEM. Asterisks denote significant effect (*p<0.05). n.s. nonsignificant. Each light gray dot corresponds to one animal.

The freezing data for mAChR blockade are presented in Figures. 6B, S4B. Baseline levels of freezing during training and test sessions did not differ between the groups (max *t*(23) = 1.30, *p* = 0.21). Fear conditioning was successful and freezing gradually increased across trials (*F*_1,23_ = 164.42, *p* < 0.001). This increase was similar across groups, as there was neither main effect of group (*F*_1,23_ = 0.084, *p* = 0.77) nor a trial by group interaction (*F*_1,23_ = 0.73, *p* = 0.40). Extinction was also successful and freezing gradually decreased across trials (*F*_1,23_ = 19.04, *p* < 0.001). Blocking mAChR with scopolamine reduced within-session freezing during extinction (*F*_1,23_ = 31.46, *p* < 0.001), and this reduction produced a trial by group interaction (*F*_1,23_ = 10.38, *p* = 0.004). However, the blockade had no effect on extinction retrieval during the post-extinction test, as all rats froze similarly (*F*_1,23_ = 0.001, *p* = 0.97). At the context tests, freezing was greater in context A than in context B (*F*_1,23_ = 41.58, *p* < 0.001), and there was no main effect of group (*F*_1,23_ = 1.39, *p* = 0.25). There was, however, a group by context interaction (*F*_1,23_ = 5.88, *p* = 0.024): vehicle controls displayed robust renewal (*F*_1,23_ = 44.72, *p* < 0.001), while the scopolamine group showed an attenuated renewal effect (*F*_1,23_ = 7.23, *p* = 0.013).

The freezing data for nAChR blockade are presented in Figs. 6D, S4D. Baseline levels of freezing during training and test sessions did not differ between the groups (max *F*_2,28_ = 2.24, *p* = 0.13). Fear conditioning was successful and freezing gradually increased across trials (*F*_1,28_ = 253.11, *p* < 0.001). This increase was similar across groups, as there was neither a main effect of group (*F*_2,28_ = 0.48, *p* = 0.62) nor a group by trial interaction (*F*_2,28_ = 1.45, *p* = 0.25). Extinction was also successful and freezing gradually decreased across trials (*F*_1,28_ = 49.26, *p* < 0.001). However, blocking nAChR in the IL reduced within-session freezing during extinction (*F*_2,28_ = 5.74, *p* = 0.008): both mecamylamine groups froze less than vehicle controls (1 µg/µl: *p* = 0.019; 10 µg/µl: *p* = 0.012) and did not differ from each other (*p* = 1). The blockade similarly reduced freezing during the post-extinction test (*F*_2,28_ = 6.30, *p* = 0.005), with both mecamylamine groups again lower than controls (1 µg/µl: *p* = 0.006; 10 µg/µl: *p* = 0.022) and not differing from each other (*p* = 1). The blockade also abolished fear renewal but only at the higher dose. There was a main effect of group (*F*_2,28_ = 4.30, *p* = 0.024), a main effect of context (*F*_1,28_ = 28.10, *p* < 0.001), and a group by context interaction (*F*_2,28_ = 7.92, *p* = 0.002). Vehicle controls renewed, freezing more in context A than in context B (*F*_1,28_ = 28.22, *p* < 0.001), as did the 1 µg/µl (*F*_1,28_ = 8.19, *p* = 0.008), whereas the 10 µg/µl group froze at similar levels in both contexts (*F*_1,28_ = 0.16, *p* = 0.69).

Together, these results indicate that blocking nAChR in the IL reproduces the effects of ^ACh^HDB→IL silencing, enhancing extinction retrieval and, at the higher dose, abolishing renewal. Blocking mAChR reproduced the effects only partially, reducing freezing within the extinction session and attenuating renewal without affecting extinction retrieval.

## Discussion

The present experiments examined the role of the ^ACh^HDB→IL pathway in fear regulation. Silencing the pathway during extinction immediately reduced freezing at the post-extinction test and, critically, abolished renewal whether it was tested in the conditioning context or in another familiar context. In two of the three experiments, silencing also reduced freezing within the extinction session itself. Ex vivo electrophysiology suggested that these effects may arise from the loss of inhibitory influence that IL interneurons exert over local projection neurons. Both ACh receptor classes contributed to this influence, with nAChRs supporting the excitation of interneurons and mAChRs required for the inhibition of projection neurons. Our pharmacological data points to a central role of nAChRs, as their blockade in the IL reproduced the effects of pathway silencing, whereas mAChR blockade attenuated renewal without affecting extinction retrieval. To the best of our knowledge, these findings are the first demonstrations of a role for the HDB→IL cholinergic pathway in fear regulation.

The main behavioral effects of our optogenetic manipulations were selective in two respects. First, they emerged only when the ^ACh^HDB→IL pathway was silenced during extinction; silencing the same pathway during conditioning had no effect. Second, silencing the ^ACh^HDB→PL pathway did not alter extinction, its retrieval, or renewal, whether applied during conditioning or extinction. This spatial and temporal specificity is consistent with the broader literature on the mPFC contribution to fear regulation (Vidal-Gonzalez et al., 2006; Corcoran and Quirk, 2007; Laurent and Westbrook, 2009; Sierra-Mercado et al., 2011; Do-Monte et al., 2015; Kim et al., 2016; Lingawi et al., 2017b; Lay et al., 2020). The ^ACh^HDB→IL results parallel our previous work (Crimmins et al., 2023), in which silencing HDB cholinergic input to the basolateral amygdala (BLA) during extinction likewise facilitated extinction retrieval and prevented renewal. Whether the two pathways serve the same function or regulate separate aspects of extinction remains an important open question, as does whether they arise from the same HDB neurons: anatomical evidence points both to populations that innervate multiple forebrain targets and to divergent, target specific ones (Barabás et al., 2024). Future work using dual labeling or intersectional approaches will be needed to resolve this. Regardless, the present study provides evidence that basal forebrain cholinergic signaling regulates fear through the IL. This conclusion aligns with recent work showing that basal forebrain territories are recruited by the IL during extinction (Fernandes-Henriques et al., 2025). Together, these findings situate the IL within a broader basal forebrain network that shapes the formation and expression of extinction memories.

The behavioral results suggest that silencing the ^ACh^HDB→IL pathway increases the activity of IL pyramidal neurons. Stimulating these neurons during extinction training immediately reduces freezing and lowers freezing during subsequent manipulation-free test (Do-Monte et al., 2015; Kim et al., 2016; Raij et al., 2018; Bayer et al., 2024), a pattern we observed in most of our experiments with pathway silencing. This interpretation aligns with our electrophysiology, which showed that HDB cholinergic input predominantly excites local interneurons, cells that may in turn impose feedforward inhibition on IL pyramidal neurons. Most of the excited interneurons we recorded were in superficial layers and resembled neurogliaform and classic-accommodating subtypes, consistent with the strong cholinergic bias toward superficial IL reported anatomically (Bloem et al., 2014b). These cells are well placed to provide the feedforward inhibition that constrains pyramidal output during extinction, although this disynaptic route is unlikely to be the whole story. Indeed, the excitation of interneurons was only partly reduced by nAChR blockade, whereas the inhibition of pyramidal neurons was abolished by mAChR, suggesting that the latter receptors probably contribute as well, either on pyramidal neurons directly or on the interneurons that inhibit them. More broadly, IL interneurons play distinct roles in fear and extinction with parvalbumin and somatostatin populations contributing differentially (Binette et al., 2023; Campos Cardoso et al., 2026). Our findings add to this picture by identifying a cholinergic route through which superficial layer interneurons may gate IL output, such that removing the cholinergic input disinhibits pyramidal neurons and may strengthen the extinction memory.

Our electrophysiology suggested that nAChR-mediated cholinergic excitation of IL interneurons accounts for part of the cholinergic influence on the IL, while a muscarinic component was required for the inhibition of pyramidal neurons. Our intracranial pharmacology revealed a behavioral dissociation between the two receptor classes: nicotinic blockade in the IL reproduced the effects of ^ACh^HDB→IL silencing, enhancing extinction retrieval and abolishing renewal, whereas muscarinic blockade reduced freezing within the extinction session and attenuated, but did not, abolish renewal, without affecting extinction retrieval. This muscarinic result contrasts with previous work (Santini et al., 2012), in which intra-IL scopolamine impaired extinction retrieval the following day. The discrepancy may reflect differences in protocol, including the lower dose of scopolamine used here, and the stronger extinction achieved by our animals, which left little room to detect a consolidation deficit. Our nicotinic results also differ from our previous findings (Crimmins et al., 2023), where systemic nAChR blockade during extinction had no effect, although the systemic route complicates direct comparison and may obscure effect localized to the IL. Regardless, the finding that the nicotinic manipulation was sufficient to render extinction context-independent points to the IL as a tractable target, and to selective, orally available nicotinic compounds such as the α7 nAChR negative allosteric modulator BNC210 (O’Connor et al., 2024) as a promising avenue.

Some aspects of our findings warrant further comment. Silencing the ^ACh^HDB→IL pathway during extinction reduced within-session freezing in two AAB experiments, in which extinction took place in the conditioning context, but not in the ABA experiment, in which extinction took place in a separate context. The source of this difference remains unclear, an explanatory analysis restricted to the first half of extinction trials revealed a significant reduction of freezing in the ABA experiment (F_(1,26)_=4.53, p=0.043). The effect of silencing on within-session freezing therefore appears to be present across experiments, but more readily detected when extinction occurs in the conditioning context and overall levels of freezing are higher. A second consideration is that the post-extinction test, here and in our previous work (Crimmins et al., 2023) may function as an additional extinction session and so confounds the assessment of renewal. This seems unlikely. The renewal test of interest is conducted in the context in which extinction did not occur (context A for ABA and context B for AAB), so any further reduction in freezing to the extinction context would increase rather than obscure the difference between contexts, and would therefore facilitate rather than hinder the detection of renewal. Finally, consistent with our previous study (Crimmins et al., 2023), we observed no sex differences in any experiment. This null should be interpreted with caution, as our experiments may have lacked power to detect such effects and we did not track the estrous cycle, despite evidence that sex hormones influence fear extinction (Pestana et al., 2026). A sex difference remains plausible because estrogen receptors are expressed by basal forebrain cholinergic neurons in a regionally asymmetric manner, being more prevalent in the HDB than in other cholinergic territories (Gibbs, 1996; Shughrue et al., 2000; Daniel and Dohanich, 2001; Gibbs, 2010). The ^ACh^HDB→IL pathway studied here may therefore be sensitive to hormonal state, making cycle stratified replication an important direction for future work.

In conclusion, we identify the ^ACh^HDB→IL pathway as a determinant of whether extinction memories remain context-specific and prone to relapse or become context-general and resistant to it. During extinction, this cholinergic input appears to engage nicotinic signaling in the IL to constrain pyramidal output and bias extinction toward context-specific memory. Removing this input, optogenetically or through nicotinic blockade, lifts the constraint, strengthening the extinction memory and abolishing renewal. These findings point to nicotinic signaling in the IL as a promising target for improving extinction-based therapies.

## Supporting information

Supplemental results

## AUTHOR CONTRIBUTIONS

BPPL and VL designed the experiments. BPPL and BCC performed the experiments and analyzed the behavioral and neural data. BPPL, SM, and VL wrote and edited the paper.

## ACKNOWLEDGEMENTS

This work was supported by the Australian Research Council in the form of a Future Fellowship (FT220100474) to VL and a Discovery Early Career Researcher Award (DE240100614) to BPPL.

## CONFLICT OF INTEREST

The authors declare no competing financial interests.

## Notes

### Competing Interest Statement

The authors have declared no competing interest.

### Summary of Updates

Manuscript modified to improve clarity, presentations and to provide conclusions better aligned with the findings.

