## Supplemental results for "Silencing basal forebrain cholinergic input to the infralimbic cortex renders fear extinction resistant to renewal"

### Supplementary Material

#### Figures and legends

#### Figure S1 – Histological and behavioral data related to Figure 2

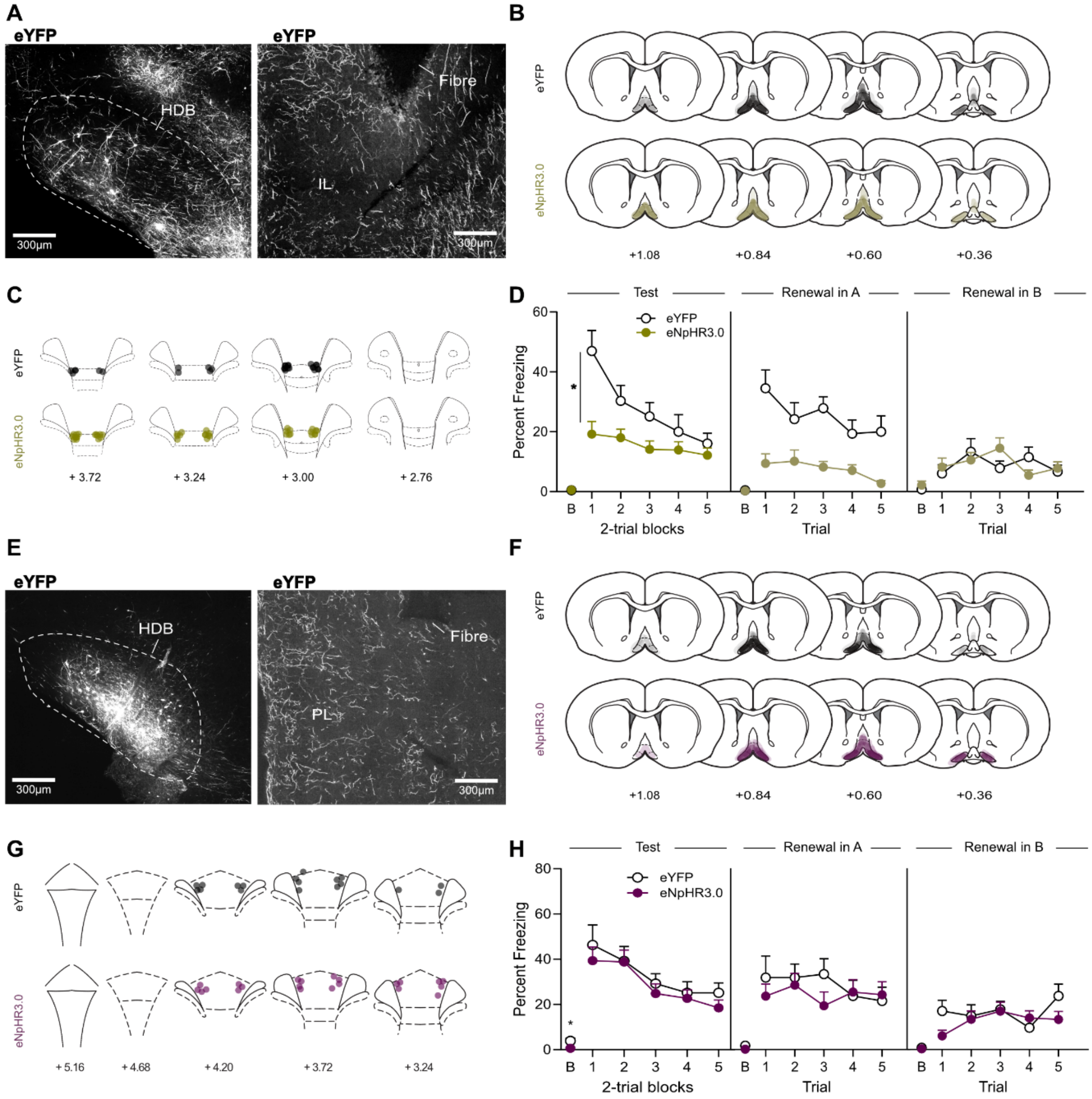


**A.** Micrographs showing eYFP viral expression in HDB cholinergic neurons (left) and eYFP-positive IL cholinergic terminals and fiber optic placements (right). **B.** Minimal (light black for eYFP and light gold for eNpHR3.0-eYFP) and maximal (darker black for eYFP and darker gold for eNpHR3.0-eYFP) extent of the HDB viral infection. Distances are indicated in mm from bregma. **C.** Location of fiber optics in the IL (black for eYFP and gold for eNpHR3.0-eYFP). Distances are indicated in mm from bregma. **D.** Baseline freezing (B) to the context prior to first tone CS presentation. $^{ACh}$HDB→IL silencing during fear extinction reduced freezing at a faster rate during the post-extinction test relative to the eYFP-control rats ($F_{1,26}$=10.50, p=0.003). At the renewal tests, freezing was higher in the conditioning context A than in the extinction context B in the control rats ($F_{1,26}$=47.47, p<0.001). By contrast, $^{ACh}$HDB→IL silencing during fear extinction abolished renewal. Rats that received $^{ACh}$HDB→IL silencing during fear extinction froze very little in both contexts ($F_{1,26}$=0.92, p=0.35). **E.** Micrographs showing eYFP viral expression in HDB cholinergic neurons (left) and eYFP-positive PL cholinergic terminals and fiber optic placements (right). **F.** Minimal (light black for eYFP and light purple for eNpHR3.0-eYFP) and maximal (darker black for eYFP and darker purple for eNpHR3.0-eYFP) extent of the HDB viral infection. Distances are indicated in mm from bregma. **G.** Location of fiber optics in the PL (black for eYFP and purple for eNpHR3.0-eYFP). Distances are indicated in mm from bregma. **H.** $^{ACh}$HDB→PL silencing during fear extinction had no effect on freezing during the post-extinction test. The rate of decline in freezing was similar between groups ($F_{1,18}$=0.007, p=0.94). At the renewal tests, all rats froze more in context A than in context B ($F_{1,18}$=15.90, p<0.001) and fear renewal was unaffected by $^{ACh}$HDB→PL silencing during fear extinction. Data are shown as mean + SEM.

**Figure S2 – Histological and behavioral data related to Figure 3**

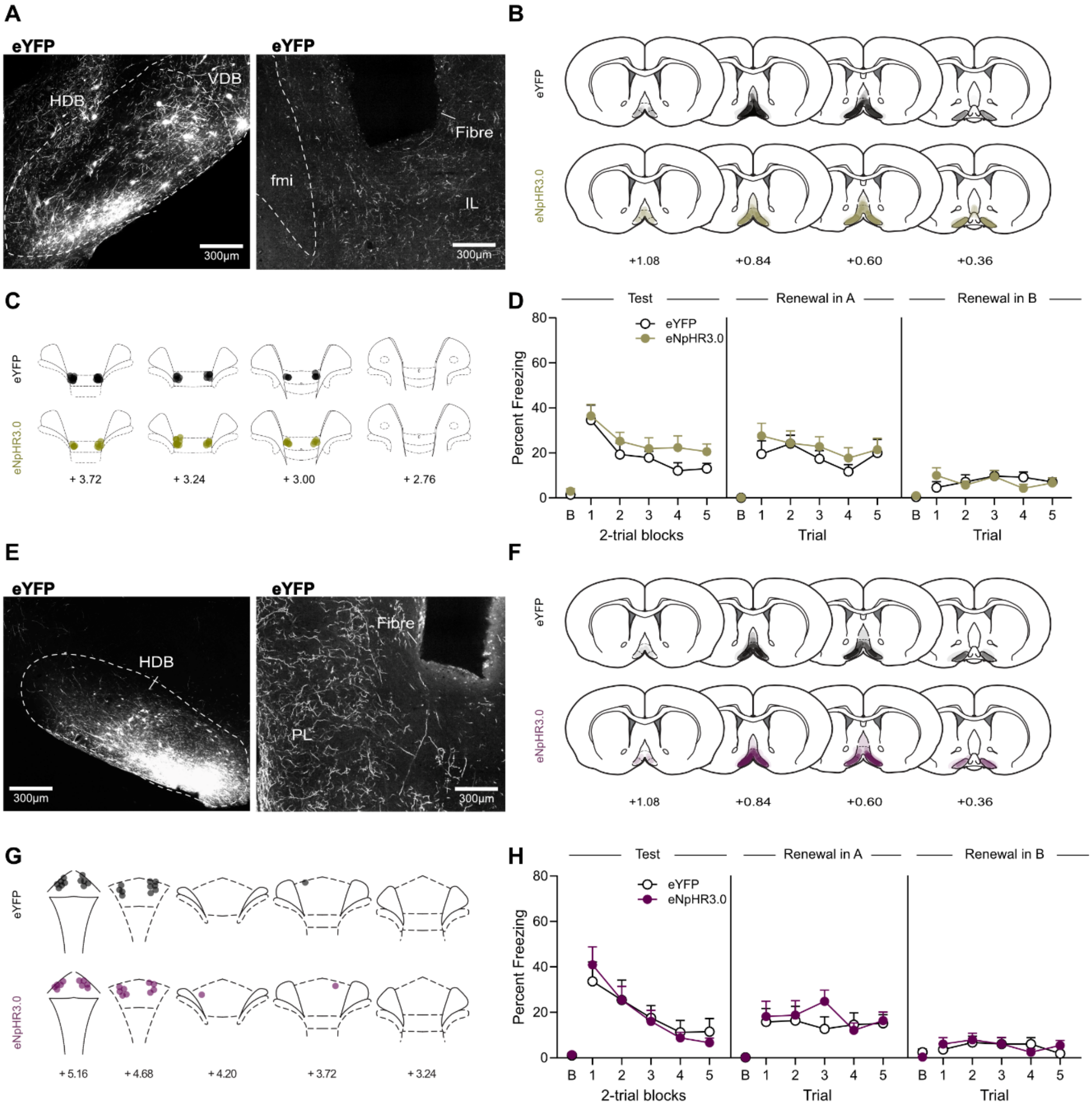


**A.** Micrographs showing eYFP viral expression in HDB cholinergic neurons (left) and eYFP-positive IL cholinergic terminals and fiber optic placements (right). **B.** Minimal (light black for eYFP and light gold for eNpHR3.0-eYFP) and maximal (darker black for eYFP and darker gold for eNpHR3.0-eYFP) extent of the HDB viral infection. Distances are indicated in mm from bregma. **C.** Location of fiber optics in the IL (black for eYFP and gold for eNpHR3.0-

eYFP). Distances are indicated in mm from bregma. **D.** Baseline freezing (B) to the context prior to first tone CS presentation. $^{ACh}$HDB→IL silencing during fear conditioning had no effect on freezing performance during the post-extinction test. The rate of decline in freezing was similar between groups ($F_{1,25}$=1.13, p=0.30). At the renewal tests, all rats froze more in context A than in context B ($F_{1,25}$=45.23, p<0.001) and fear renewal was unaffected by $^{ACh}$HDB→IL silencing during fear conditioning. **E.** Micrographs showing eYFP viral expression in HDB cholinergic neurons (left) and eYFP-positive PL cholinergic terminals and fiber optic placements (right). **F.** Minimal (light black for eYFP and light purple for eNpHR3.0-eYFP) and maximal (darker black for eYFP and darker purple for eNpHR3.0-eYFP) extent of the HDB viral infection. Distances are indicated in mm from bregma. **G.** Location of fiber optics in the PL (black for eYFP and purple for eNpHR3.0-eYFP). Distances are indicated in mm from bregma. **H.** $^{ACh}$HDB→PL silencing during fear conditioning had no effect on freezing performance during the post-extinction test. The rate of decline in freezing was similar between groups ($F_{1,20}$=1.30, p=0.27). At the renewal tests, all rats froze more in context A than in context B ($F_{1,20}$=22.84, p<0.001) and fear renewal was unaffected by $^{ACh}$HDB→PL silencing during fear conditioning. Data are shown as mean + SEM.

**Figure S3 – Histological and behavioral data related to Figure 4**

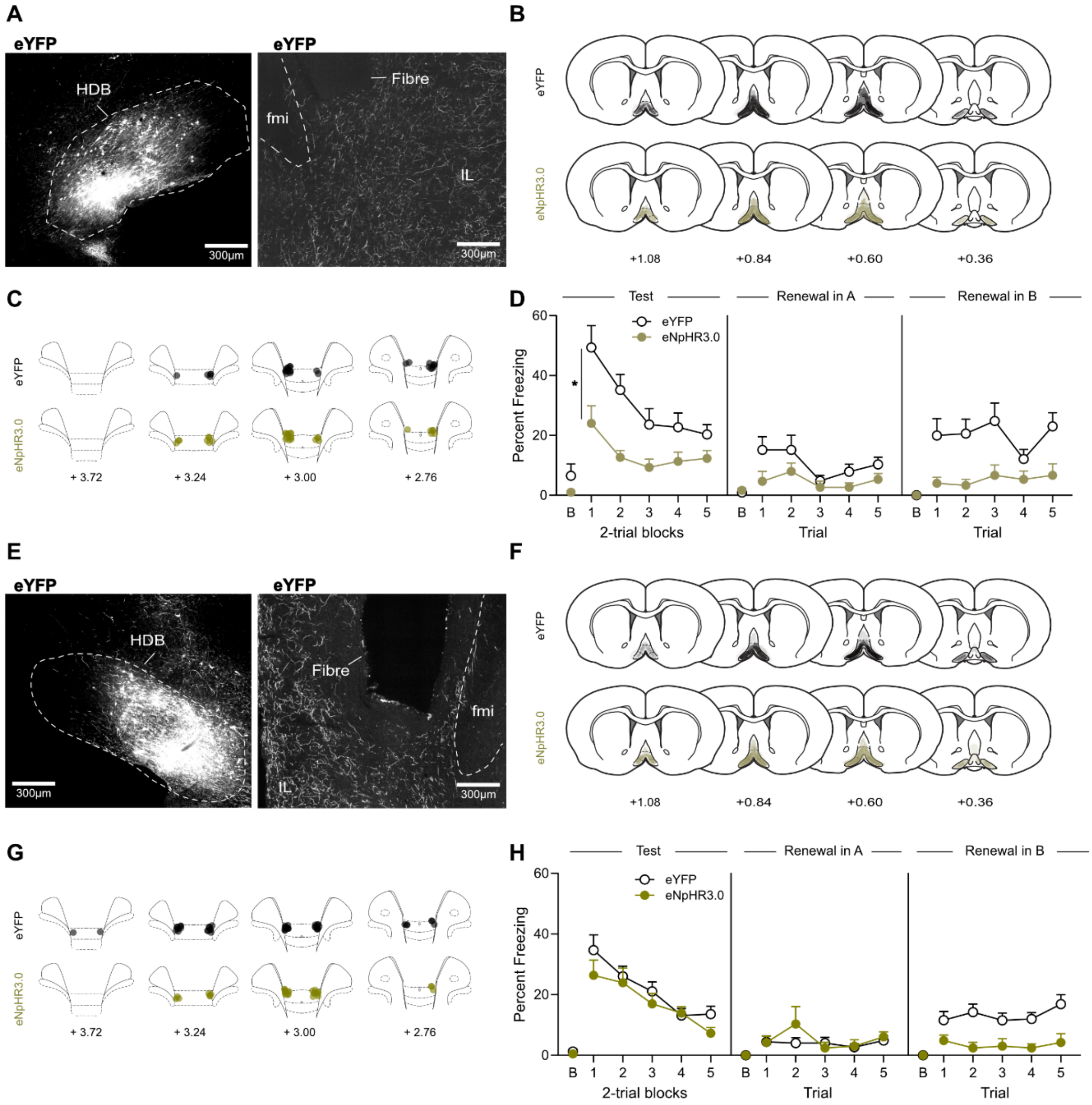


**A.** Micrographs showing eYFP viral expression in HDB cholinergic neurons (left) and eYFP-positive IL cholinergic terminals and fiber optic placements (right). **B.** Minimal (light black for eYFP and light gold for eNpHR3.0-eYFP) and maximal (darker black for eYFP and darker gold for eNpHR3.0-eYFP) extent of the HDB viral infection. Distances are indicated in mm from bregma. **C.** Location of fiber optics in the IL (black for eYFP and gold for eNpHR3.0-eYFP). Distances are indicated in mm from bregma. **D.** Baseline freezing (B) to the context prior to first tone CS presentation. $^{Ach}$HDB→IL silencing during fear extinction reduced freezing at a faster rate during the post-extinction test relative to the eYFP-control rats ($F_{1,19}=6.27$, $p=0.022$). At the renewal tests, freezing was higher in the conditioning context A than in the novel context B in the control rats ($F_{1,19}=18.59$, $p<0.001$). By contrast, $^{ACh}$HDB→IL silencing during fear extinction abolished renewal. Rats that received $^{ACh}$HDB→IL silencing during fear extinction froze very little in both contexts ($F_{1,19}=0.054$, $p=0.82$). **E.** Micrographs showing eYFP viral expression in HDB cholinergic neurons (left) and eYFP-positive IL cholinergic terminals and fiber optic placements (right). **F.** Minimal (light black for eYFP and light gold for eNpHR3.0-eYFP) and maximal (darker black for eYFP and darker gold for eNpHR3.0-eYFP) extent of the HDB viral infection. Distances are indicated in mm from bregma. **G.** Location of fiber optics in the IL (black for eYFP and gold for eNpHR3.0-eYFP). Distances are indicated in mm from bregma. **H.** $^{ACh}$HDB→IL silencing during fear extinction had no effect on freezing during the post-extinction test. The rate of decline in freezing was similar between groups ($F_{1,24}$=0.17, p=0.68). Notably, at the renewal tests, freezing was higher in the conditioning context A than in the novel context B in the control rats ($F_{1,24}$=20.32, p<0.001). By contrast, $^{ACh}$HDB→IL silencing during fear extinction abolished renewal. Rats that received $^{ACh}$HDB→IL silencing during fear extinction froze very little in both contexts ($F_{1,24}$=0.58, p=0.46). Data are shown as mean + SEM.

**Figure S4 – Histological and behavioral data related to Figure 6**

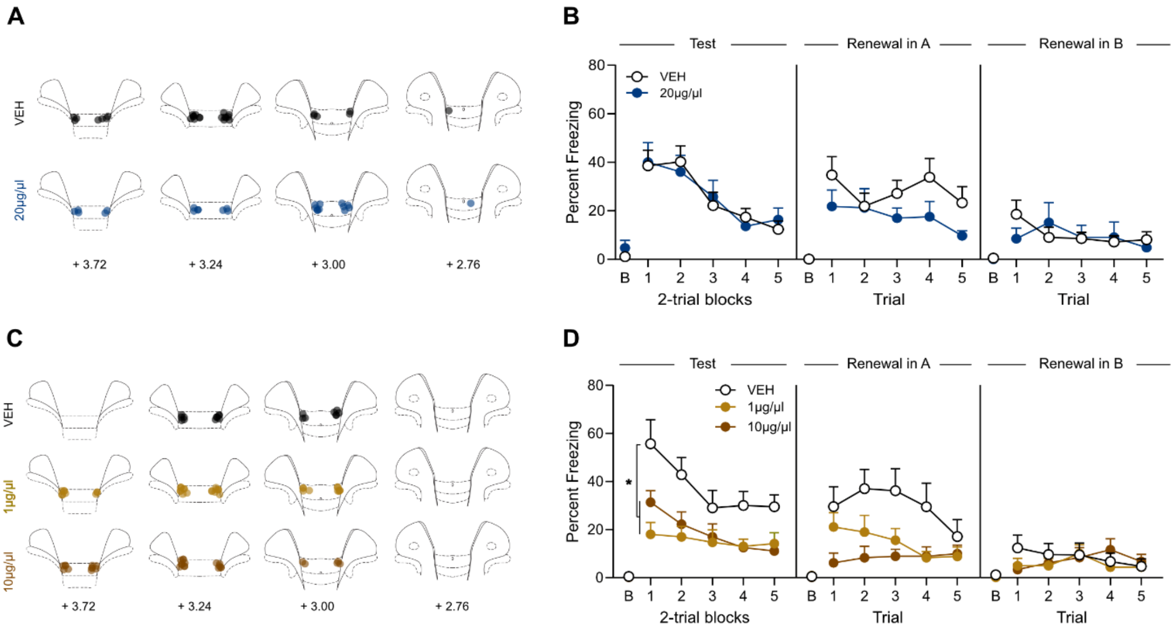


**A.** Location of cannula placements in the IL (black for vehicle and blue for 20 µg/µl). Distances are indicated in mm from bregma. **B.** Muscarinic receptor blockade during fear extinction had no effect on freezing during the post-extinction test. The rate of decline in freezing was similar between groups ($F_{1,23}$=0.06, p=0.81). **C.** Location of cannula placements in the IL (black for vehicle, light brown for 1 µg/µl, and dark brown for 10 µg/µl). Distances are indicated in mm from bregma. **D.** Baseline freezing (B) to the context prior to first tone CS presentation. Nicotinic receptor blockade during fear extinction reduced freezing at a faster rate during the post-extinction test relative to the vehicle-controls ($F_{2,28}$=6.19, p=0.006). Data are shown as mean + SEM.
